# Dopamine D2 receptors modulate intrinsic properties and synaptic transmission of parvalbumin interneurons in the mouse primary motor cortex

**DOI:** 10.1101/802140

**Authors:** Jérémy Cousineau, Léa Lescouzères, Anne Taupignon, Lorena Delgado-Zabalza, Emmanuel Valjent, Jérôme Baufreton, Morgane Le Bon-Jégo

## Abstract

Dopamine (DA) plays a crucial role in the control of motor and higher cognitive functions such as learning, working memory and decision making. The primary motor cortex (M1), which is essential for motor control and the acquisition of motor skills, receives dopaminergic inputs in its superficial and deep layers from the midbrain. However, the precise action of DA and DA receptor subtypes on the cortical microcircuits of M1 remains poorly understood. The aim of this work was to investigate how DA, through the activation of D2 receptors (D2R), modulates the cellular and synaptic activity of M1 parvalbumin-expressing interneurons (PVINs) which are crucial to regulate the spike output of pyramidal neurons (PNs). By combining immunofluorescence, *ex vivo* electrophysiology, pharmacology and optogenetics approaches, we show that D2R activation increases neuronal excitability of PVINs and GABAergic synaptic transmission between PVINs and PNs in layer V of M1. Our data reveal a mechanism through which cortical DA modulates M1 microcircuitry and might participate in the acquisition of motor skills.

**Significance Statement:** Primary motor cortex (M1), which is a region essential for motor control and the acquisition of motor skills, receives dopaminergic inputs from the midbrain. However, precise action of dopamine and its receptor subtypes on specific cell types in M1 remained poorly understood. Here, we demonstrate in M1 that dopamine D2 receptors (D2R) are present in parvalbumin interneurons (PVINs) and their activation increases the excitability of the PVINs, which are crucial to regulate the spike output of pyramidal neurons (PNs). Moreover the activation of the D2R facilitates the GABAergic synaptic transmission of those PVINs on layer V PNs. These results highlight how cortical dopamine modulates the functioning of M1 microcircuit which activity is disturbed in hypo- and hyperdopaminergic states.

## Introduction

The neuromodulator dopamine (DA) plays a key role in the ability of neural circuits to adaptively control behaviour (Schultz, 2007; Vitrac and Benoit-Marand, 2017; Berke, 2018). Indeed, the DA system plays a major role in motor and cognitive functions through its interactions with several brain regions, and its dysregulation leads to cognitive dysfunction (Duvarci et al., 2018) and pathologies like Parkinson’s disease and schizophrenia (Nieoullon, 2002). Recently, it has been suggested that the primary motor cortex (M1) may also be influenced by dopamine (Hosp and Luft, 2013; Guo et al., 2015). The architecture of the dopaminergic inputs to M1 has been well characterized anatomically. Coming mainly from the ventral tegmental area (VTA) but also from the *substantia nigra pars compacta* (SNc), they richly innervate the superficial and deep layers of the rodent and primate M1 (Descarries et al., 1987; Lewis et al., 1987; Vitrac et al., 2014; Hosp et al., 2015). However, their functional significance is poorly understood and reports of their effects remain conflicting, presumably because of the *in vivo* exploration and wide neuronal diversity in M1 (Hosp and Luft, 2013; Vitrac et al., 2014; Vitrac and Benoit-Marand, 2017).

DA acts *via* two main classes of receptors, the D1-like (D1R) and the D2-like family (D2R) which differentially modulate adenylyl cyclase (Beaulieu and Gainetdinov, 2011). In M1, both families of DA receptors are present in the deep layers (Dawson et al., 1986; Lidow et al., 1989; Weiner et al., 1991; Gaspar et al., 1995). Based on *in situ* hybridization, it appears that layer V of the cortex, the layer where pyramidal neurons (PNs) integrate inputs from many sources and distribute information to cortical and subcortical structures, mainly contains D2R mRNA (Gaspar et al., 1995). Previous work has described the effect of DA on neuronal activity in M1 neurons *in vivo*, but most of these studies focused on PNs and draw different conclusions regarding an inhibitory or excitatory effect of DA on neuronal activity in M1 (Awenowicz and Porter, 2002; Vitrac et al., 2014). However, there is a large body of evidence supporting that inhibition is important in controlling the excitatory circuits. Among the various interneurons (INs) (Anon, 2008; DeFelipe et al., 2013; Lodato et al., 2015; Markram et al., 2015), parvalbumin-expressing INs (PVINs) represent a minority cell type. However, they are crucial for normal brain function (Donato et al., 2013; Courtin et al., 2014): they powerfully regulate the spike output of PNs, mainly by targeting their somatic and perisomatic regions (Hu et al., 2014). In addition, they are also recruited for motor execution (Estebanez et al., 2017).

To better understand the cellular and network basis of DA action in M1, it is necessary to determine the cellular targets of DA innervation. We hypothesized that DA in M1 contributes to normal microcircuit processing by modulating the activity of PVINs in layer V through D2R. To test this hypothesis, we first performed qualitative mapping of the M1 neuronal population expressing D2R and electrophysiologically characterized these D2R-positive neurons. Then, we investigated the impact of D2R activation on the excitability of PVINs using patch-clamp electrophysiology and on GABAergic synaptic transmission between PVINs and PNs using optogenetics. We found that D2Rs are broadly expressed in M1, in both superficial and deep layers. In layer V, the majority of neurons expressing D2R are PVINs. Moreover, D2R agonists increase the excitability of PVINs and also enhance GABAergic synaptic transmission between PVINs and PNs. Our results clarify and highlight the role of DA in modulating the activity of cortical microcircuits in M1.

## Methods

### Animals

All experiments were performed in accordance with the guidelines of the French Agriculture and Forestry Ministry for handling animals (authorization number/license D34-172-13 and APAFIS #14255). C57BL6J and three transgenic mouse lines were used for this study. Drd2-Cre:Ribotag mice were used for the morphological study (Puighermanal et al., 2015). PV-Cre:Ai9T mice were generated by crossing PV-Cre mice (B6;129P2-*PValb*^*tm1(cre)Arbr*^/J; JAX stock # 008069; (Kaiser et al., 2016)) with Ai9T mice (B6.Cg-*Gt(ROSA)26Sor*^*tm9(CAG-tdTomato)Hze*^/J; stock # 007909) and the Drd2-Cre:Ai9T line was generated by crossing Drd2-Cre mice (Tg(Drd2-Cre)ER44Gsat; Gensat Project at Rockefeller University) with Ai9T mice (B6.Cg-*Gt(ROSA)26Sor*^*tm9(CAG-tdTomato)Hze*^/J; stock # 007909). These two lines express the red fluorescent protein double-tomato (tdTom) under endogenous regulatory elements of the parvalbumin gene locus and those of D2R, respectively. Males and females, 8-12 weeks old, were used for *ex vivo* experiments. All animals were maintained in a 12h light/dark cycle, in stable conditions of temperature and humidity, with access to food and water *ad libitum*.

### Tissue preparation and immunofluorescence

Male Drd2-Cre:Ribotag mice, 8-10 weeks old (n=6), were used for the morphological study. Mice were rapidly anesthetized with Euthasol (360 mg/kg, i.p., TVM lab, France) and transcardially perfused with 4% (w/v) paraformaldehyde in 0.1 M sodium phosphate buffer (pH 7.5). Brains were post-fixed overnight in the same solution and stored at 4°C. Sections of 30 µm were cut with a vibratome (Leica, France) and stored at −20°C in a solution containing 30% (v/v) ethylene glycol, 30% (v/v) glycerol and 0.1 M sodium phosphate buffer until they were processed for immunofluorescence. M1 motor cortex sections were identified using a mouse brain atlas; sections located between +1.60 and +0.98 mm from bregma were included in the analysis (Franklin and Paxinos, 2007). Sections were processed as follows: free-floating sections were rinsed 3 x 10 min in Tris-buffered saline (TBS, 50 mM Tris-HCl, 150 mM NaCl, pH 7.5). After 15 min incubation in 0.2% (v/v) Triton X-100 in TBS, sections were rinsed again in TBS and blocked for 1 h in a solution of 3% BSA in TBS. Finally, they were incubated 72 h at 4°C in 1% BSA, 0.15% Triton X-100 with the primary antibodies (Table 1). Sections were rinsed 3 x 10 min in TBS and incubated for 45 min with goat Cy3-coupled anti-rabbit or anti-mouse (1:500, Jackson ImmunoResearch Labs Cat# 111-165-003) and goat Alexa Fluor 488-coupled anti-mouse or anti-rabbit (1:500, Thermo Fisher Scientific Cat# A-11039). Sections were rinsed 2 x 10 min in TBS and twice in Tris-buffer (1 M, pH 7.5) before mounting in DPX (Sigma-Aldrich). Confocal microscopy and image analysis were carried out at the Montpellier RIO Imaging Facility. All images covering the M1 motor cortex were single confocal sections acquired using sequential laser scanning confocal microscopy (Leica SP8) and stitched together as a single image. Double-labeled images from each region of interest were also single confocal sections obtained using sequential laser scanning confocal microscopy (Leica SP8). HA-immunopositive cells were pseudo-colored cyan and other immunoreactive markers were pseudo-colored orange. Images used for quantification were all single confocal sections. HA-positive cells were manually counted using the cell counter plugin of the ImageJ software in M1 motor cortex, taking into account the cortical layers (layer I, layers II-III and layers V-VI). Adjacent serial sections were never counted for the same marker to avoid any double counting of hemisected neurons. Values in the histograms in Figure 1B represent the percentage of HA-expressing neurons in layer I, layer II-III and Layer V-VI (n = 5-6 mice). Total numbers of HA- and marker-positive cells counted are indicated between parentheses.

**Table 1:**
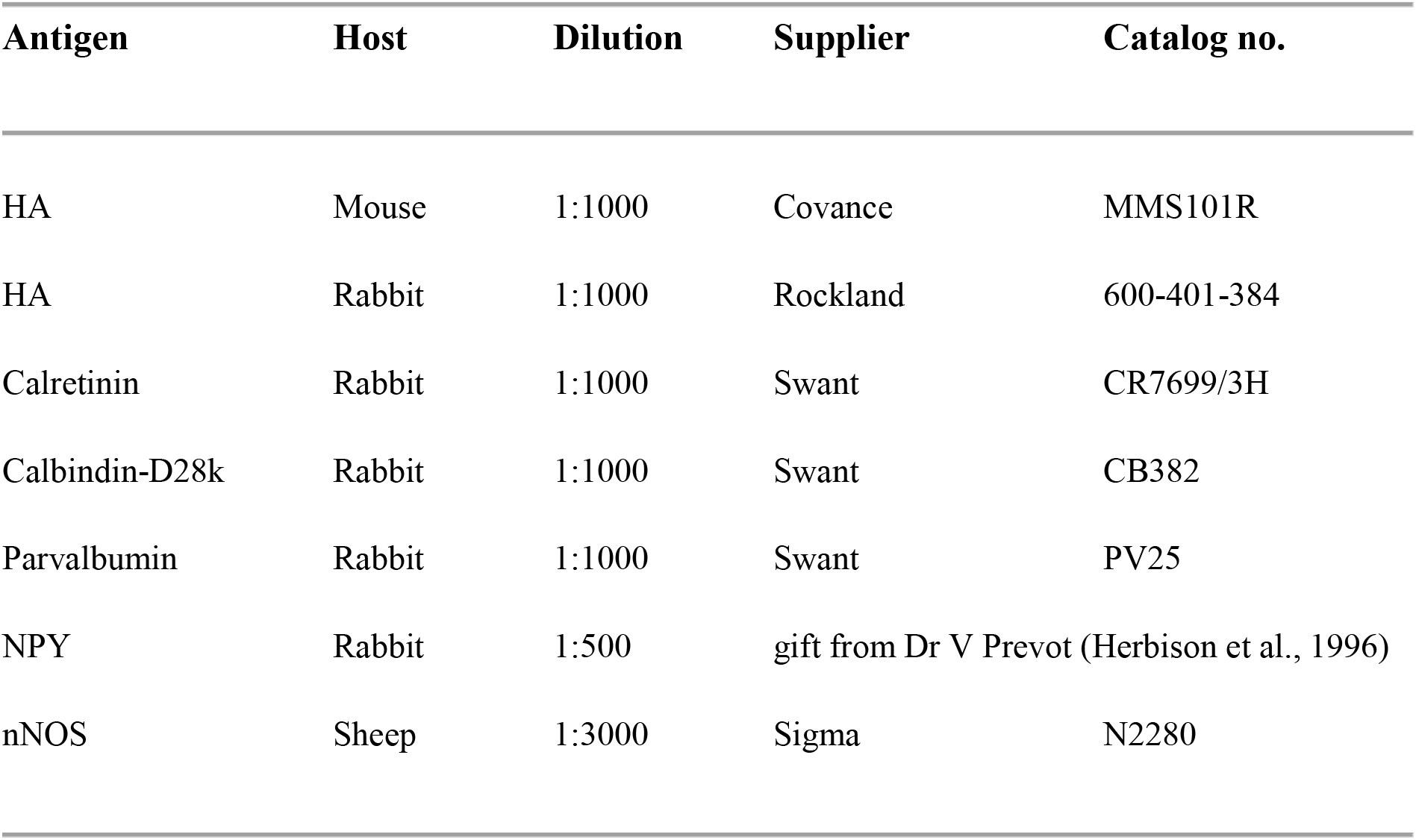
List of primary antibodies

**Figure 1:**
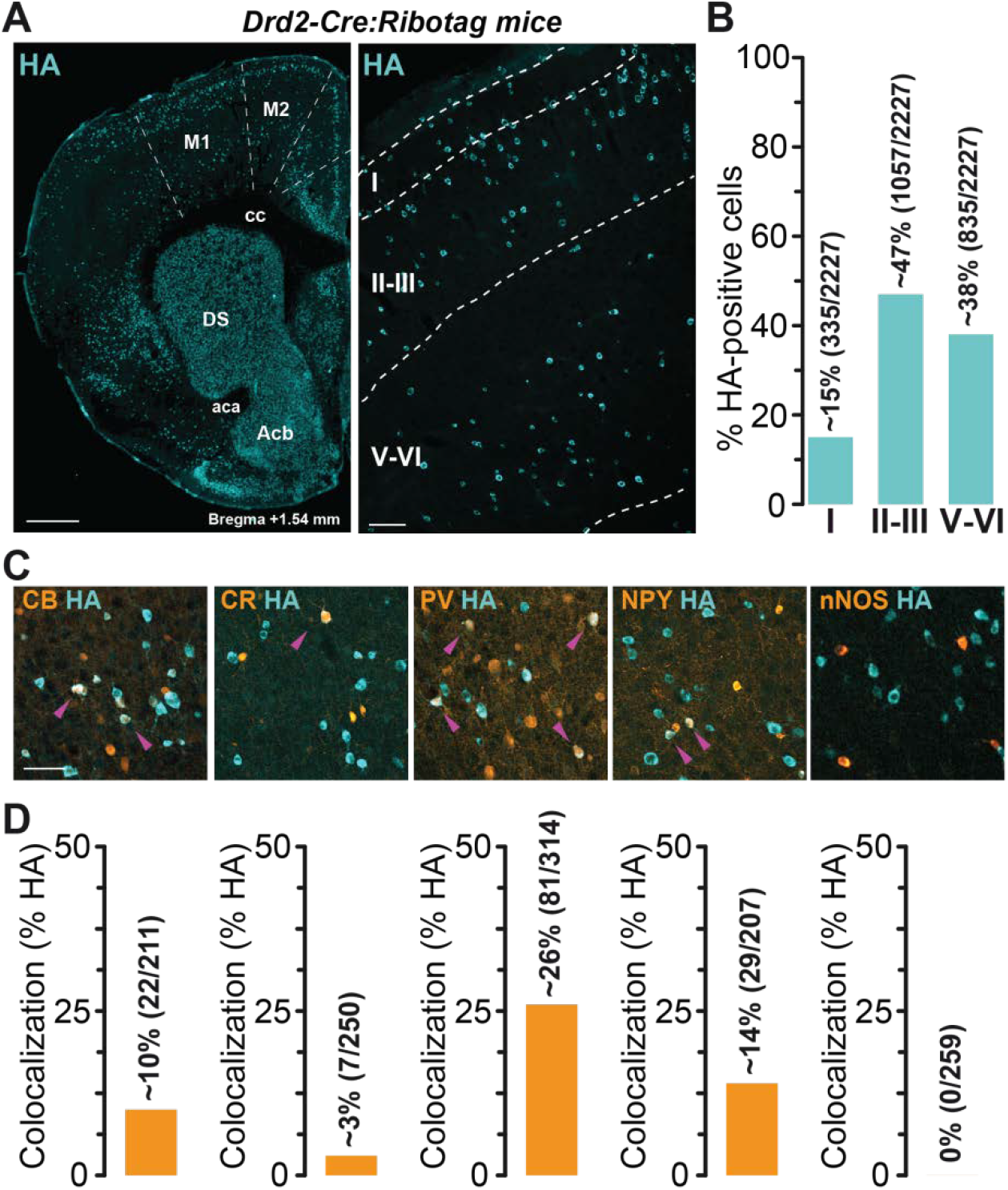
Distribution of D2R-expressing neurons in the M1 motor cortex in *Drd2-Cre:Ribotag* mice. (A) Coronal section from *Drd2-Cre:Ribotag* mice stained with HA showing the distribution of D2R-expressing neurons in the different layers of M1 motor cortex. Scale bars: 500 μm (left), 50 μm (right). (B) Histogram showing the distribution of HA-labeled neurons in layer I, layers II-III and layers V-VI of the M1 motor cortex (17 hemispheres analyzed, 5 mice). The distribution is expressed as a percentage of HA-positive neurons in all layers. The number of HA-positive cells counted is indicated between parentheses. (C) HA (cyan) and calbindin-D28k (CB), calretinin (CR), parvalbumin (PV), neuropeptide Y (NPY) and nNOS (orange) immunofluorescence in M1 motor cortex layers V-VI of *Drd2-Cre:Ribotag* mice. Magenta arrowheads indicate HA/markers-positive neurons. Scale bars: 40 μm. (D) Histograms showing the co-expression as a percentage of HA-positive cells in M1 motor cortex layers V-VI of *Drd2-Cre:Ribotag* mice. The total numbers of HA- and marker-positive cells counted are indicated between parentheses. DS: dorsal striatum; cc: corpus callosum; Acb: nucleus accumbens; aca: anterior commissure.

### Slice preparation

Coronal sections containing M1 were prepared from 8 to 12 week-old mice. Mice were first sedated by inhaling isoflurane (4%) for approximately 30 s and then deeply anesthetized with a mixture of ketamine and xylazine (100 and 20 mg/kg, i.p., respectively). After the disappearance of the reflexes, a thoracotomy was performed to allow transcardial perfusion of a saturated (95% O_2_/5% CO_2_) ice-cold solution containing 250 mM sucrose, 10 mM MgSO_4_⋅7H_2_O, 2.5 mM KCl, 1.25 mM NaH_2_PO_4_⋅H_2_O, 0.5 mM CaCl_2_⋅H_2_O, 1.3 mM MgCl_2_, 26 mM NaHCO_3_ and 10 mM D-glucose. After decapitation, each brain was quickly removed and cut into coronal slices (300-350 µm) using a vibratome (VT-1200S; Leica Microsystems, Germany). The slices were then incubated at 34°C for 1 h in a standard artificial cerebrospinal fluid (ACSF) saturated by bubbling 95% O_2_/5% CO_2_ and containing 126 mM NaCl, 2.5 mM KCl, 1.25 mM NaH_2_PO_4_⋅H_2_O, 2 mM CaCl_2_⋅H_2_O, 2 mM MgSO_4_⋅7H_2_O, 26 mM NaHCO_3_ and 10 mM D-glucose, supplemented with 5 µM glutathion and 1 mM sodium pyruvate. Slices were maintained at room temperature in the same solution until recording.

### Electrophysiology

Whole-cell patch-clamp experiments were performed in a submersion recording chamber under an upright microscope (Ni-E workstation, Nikon). Slices were bathed in ACSF containing 126 mM NaCl, 3 mM KCl, 1.25 mM NaH_2_PO_4_⋅H_2_O, 1.6 mM CaCl_2_⋅H_2_O, 2 mM MgSO_4_⋅7H_2_O, 26 mM NaHCO_3_ and 10 mM D-glucose. M1 layer V neurons were visualized with infrared differential interference contrast and fluorescence microscopy (Spectra X light engine, Lumencor) (Froux et al., 2018). Pyramidal neurons (PNs) were identified on morphological criteria (triangle-shaped soma) and D2R-positive cells and PV-positive interneurons (PVINs) were identified by the fluorescence of tdTom. Recording electrodes were pulled from borosilicate glass capillaries (G150–4; Warner Instruments, Hamden, CT, USA) with a puller (Sutter Instrument, Model P-97) and had a resistance of 5-7 MΩ. They contained 135 mM K-Gluconate, 3.8 mM NaCl, 1 mM MgCl_2_⋅6H_2_O, 10 mM Hepes, 0.1 mM Na_4_EGTA, 0.4 mM Na_2_GTP and 2 mM MgATP for the current-clamp experiments. For the recordings of spontaneous and miniature inhibitory post-synaptic currents (sIPSCs and mIPSCs, respectively) in voltage clamp experiments, K-Gluconate was replaced by CsCl and 2 mM Qx-314 was added to prevent action potentials. In all cases, the osmolarity of the intrapipette solution was between 285 and 295 mOsm and pH was adjusted to 7.2. Experiments were conducted using a Multiclamp 700B amplifier and Digidata 1440 digitizer controlled by Clampex 10.3 (Molecular Devices, Sunnyvale, CA, USA) at 34°C. Data were acquired at 20 kHz and low-pass filtered at 4 kHz. Whole-cell patch clamp recordings with CsCl- or K-Glu-filled electrodes were corrected for a junction potential of 4 mV and 13 mV, respectively. In voltage clamp experiments, series resistance was continuously monitored by a step of −5 mV. Data were discarded when the series resistance increased by >20%. sIPSCs and mIPSCs were recorded at a holding potential of −64 mV.

To evaluate their intrinsic excitability, neurons were injected with increasing depolarizing current pulses (50 pA steps, ranging from 0 to +550 pA, 1000 ms duration). Action potential firing frequency was calculated for each current pulse. To measure the input resistance, a hyperpolarizing −100 pA pulse current of 1 s was applied and the voltage response was measured at steady state. Input-output curves (F-I curves, frequency of action potential firing as a function of injected current) were constructed.

### Drugs

Unless otherwise stated, drugs were prepared in distilled water as concentrated stock solutions and stored at −20°C. Drugs were diluted daily at the experimental concentrations and perfused in the recording chamber. When indicated, ionotropic glutamatergic and GABAergic transmissions were blocked. NMDA receptors were inhibited by 50 µM D-(-)-2-amino-5-phosphonopentanoic acid (APV); AMPA/kainate receptors by 20 µM 6,7-dinitroquinoxaline-2,3-dione (DNQX); and GABA_A_ receptors by 50 µM picrotoxin. To study spontaneous or evoked inhibitory post-synaptic currents, glutamate and GABA_B_ receptors were blocked by APV, DNQX and 1 µM (2S)-3-[[(1S)-1-(1,4-dichlorophenyl)ethyl]amino-2 hydroxypropyl](phenylmethyl) phosphinic acid (CGP 55845, dissolved in DMSO). The D2-like dopamine receptor agonist (4aR-trans)-4,4a,5,6,7,8,8a,9-octahydro-5-propyl-1H-pyrazolo[3,4-g]quino-line hydrochloride (quinpirole, 2µM) and antagonist (sulpiride, 2µM) were used. Sulpiride was dissolved in dimethylsulfoxide (DMSO). Drug effects were measured at least 10 min after drug perfusion. Chemicals were purchased from Tocris Bioscience (UK), Abcam (France) or Sigma-Aldrich (France).

### Optogenetics

To specifically activate PVINs, the cation channelrhodopsin-2 (ChR2) was expressed in PVINs within M1. To this end, the viral vector AAV2.5-EF1a-DIO-hChR2(H134R)-EYFP.WPRE.hGH (V2109TI; 6.72e^12^ gc/mL; UNC Vector Core) was injected in M1 of PV-Cre:Ai9T mice. 10 mice received 3 unilateral injections of 0.5 µL viral vector solution in M1 at the following stereotaxic coordinates (from bregma): lateral, 1.125/1.125/1.375 mm, posterior, +1.4/+1.15/+1.4 mm and depth, −1.275/-1.275/-1.475 mm. The viral vector was pressure-injected using a picospritzer III (Intracel) connected to a glass pipette at a rate of 100 µL/min. After the injection, the pipette was left in place for 1 min before being slowly retracted. Animal were housed for 2-3 weeks before electrophysiological recordings. A LED-light source (473 nm, 100 mW; Prizmatix Ltd.) was connected to an optic fiber (Ø: 500 µm; numeric aperture: 0.63) placed close to the region of interest. Single or 10 Hz trains of light pulses of 1 ms duration were used to evoke synaptic transmission between PVIN-expressing ChR2 and PNs.

### Experimental design and statistical analysis

Data analyses were performed with the Clampfit routine, Origin 7 and a custom-made software for the detection and measurement of sIPSCs and mIPSCs (Detection Mini 8.0). Statistical analysis was performed with Prism 5 (GraphPad Software, La Jolla, CA). Population data are presented as mean ± SEM. Paired data were compared using the Wilcoxon signed rank (WSR) test. Comparisons of F-I relationships were performed with a two-way repeated-measures ANOVA test followed by a Bonferroni test for multiple comparisons (Bichler et al., 2017). The Kolmogorov-Smirnov test (KS) was used to compare the cumulative distributions. Data were considered statistically significant at p < 0.05 (*p < 0.05, **p < 0.01, ***p < 0.001, n.s. not significant).

## Results

### Distribution of D2R-expressing cells in the M1 motor cortex of Drd2-Cre:Ribotag mice

We took advantage of the *Drd2-Cre:Ribotag* mice (Puighermanal et al., 2015), which express ribosomal protein Rpl22 tagged with the hemagglutinin (HA) epitope selectively in D2R-positive cells, to determine the expression pattern of D2R-positive cells in the M1 motor cortex. The analysis of HA-immunoreactivity revealed that D2R-expressing cells are distributed in all cortical layers, with the highest density in layers II-III (~47%), followed by the deep layers (V-VI) (~38%) and layer I (~15%) (Figure 1A, B). To determine the molecular identity of D2R-expressing cells located in layers V-VI of the M1 motor cortex, we performed double immunostaining and quantified the degree of co-localization of HA-immunoreactive neurons with markers of distinct classes of interneurons (Figure 1C, D) (Ascoli et al., 2008). As illustrated in Figure 1C and quantified in Figure 1D, HA-positive cells mainly correspond to PV-containing interneurons (~26%) and to a lesser extent, Calbindin-D28k (CB)- and Neuropeptide Y (NPY)-positive interneurons (~10% and 14%). In contrast, Calretinin (CR)/HA co-labeled cells represent only ~3% of HA-positive cells, while neuronal NO synthase (nNOS)/HA neurons were not detected. Although D2R-positive neurons of layer V-VI might constitute a subpopulation of cortical interneurons, our results revealed that they largely correspond to PV interneurons.

### Electrophysiological characterization of M1 D2R-expressing cells

To determine the intrinsic properties of layer V M1 D2R neurons, whole-cell patch-clamp recordings were performed using *ex vivo* slices from Drd2-Cre:Ai9T (Figure 2). We patched neurons in acute brain slices and among the 19 neurons we recorded, found 3 types of D2R-positive neurons differing in their electrophysiological properties and the shape of their soma (Figure 2). 55% were fast spiking interneurons (FS), 30% were regular spiking non-pyramidal (RSNP) and 15% were PNs. The FS neurons had a mean resting potential of −83.86 ± 2.02 mV (n=11) and were able to fire fast action potentials at a high constant rate. Their action potentials had a short duration and a large afterhyperpolarization (AHP) (Figure 2B, inset) which are general characteristics of FS neurons. The discharge frequency increased as a function of the stimulation intensity and the maximal frequency, measured for high intensities of depolarizing currents ranging from 100 to 230 Hz (Figure 2D). Their rheobase differed from one neuron to another and were on average 154.5 ± 17.13 pA. In addition, FS cells had a small input resistance (between 80 to 200 MΩ, except for a neuron). We performed immunohistochemistry to detect the expression of PV in 7 neurons filled with biocytin during whole-cell recording. 6 of 7 were PV-immunoreactive (not shown).

**Figure 2:**
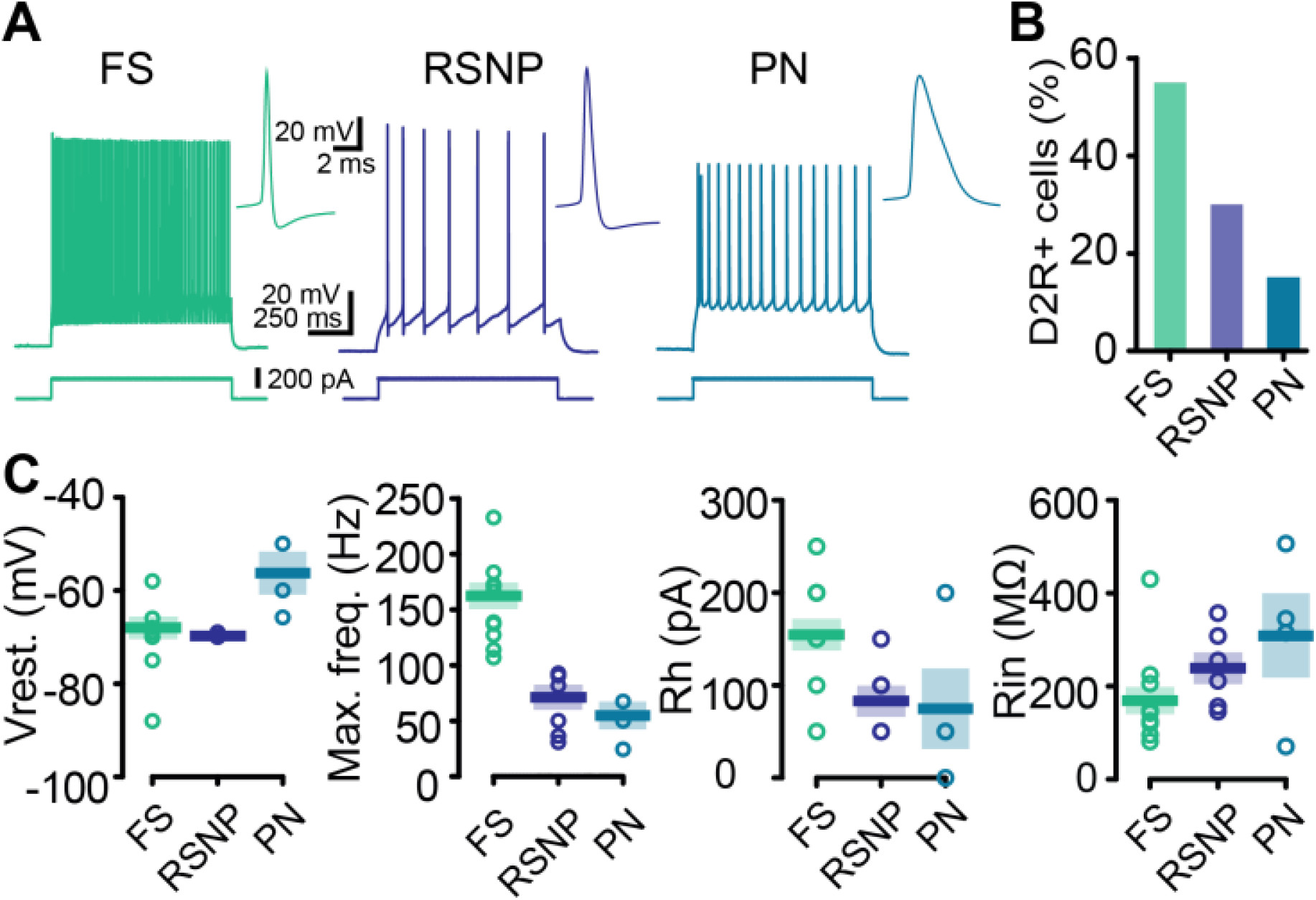
Electrophysiological characterization of D2R-expressing neurons in motor cortex M1 in *Drd2-Cre:Ai9T* mice. (A) Firing behavior of the 3 types of D2R-expressing neurons in layer V of M1. A depolarizing current injection (200 pA, 1 s) evoked a high frequency spike firing pattern in FS (left) and a lower frequency of discharge in RSNP (middle) and PNs (right). Next to each trace, an expanded view of single spikes and afterhyperpolarization is presented for the 3 groups of neurons. (B) Histogram showing the percentage of each type of D2R-expressing neurons in layer V of M1 (n = 21). (C) Summary of resting membrane potential (Vrest.), maximal firing frequency (Max. freq.), rheobase (Rh) and input resistance (Rin) in the 3 cell types.

The second cell type did not maintain high frequency repetitive discharges and was classified as RSNP because of the shape of the soma (Figure 2B, central panel). Action potentials evoked by current injection in RSNP cells had a longer duration and a relatively smaller AHP than those recorded in FS cells. All RSNP cells displayed a resting potential close to −81.23 ± 0.93 mV (n = 6). They had a low maximal frequency of discharge associated with a low rheobase. At a low discharge frequency, RSNP cells emitted action potentials with moderate or no accommodation. Finally, a few PNs were identified by the triangular shape of their soma. They exhibited a sustained action potential discharge in response to depolarizing current pulses with a low maximal frequency of discharge (Figure 2C). PNs had a mean resting potential of −71.67 ± 6.45 mV, a mean input resistance of 309.5 ± 89.97 MΩ and a mean rheobase of 75.00 ± 43.30 pA (n = 4).

### D2R activation increases the intrinsic excitability of PVINs

Since the majority of D2R cells recorded in layer V of M1 were FS interneurons and expressed PV, we switched to the PV-Cre:Ai9T mouse line to focus on the parvalbumin-expressing interneurons (PVINs), which are also mainly FS interneurons (Hu et al., 2014). In PV-Cre:Ai9T brains, PVINs can be easily targeted for recording as they express the fluorescent protein tdTom. We investigated the effect of a typical D2 agonist, quinpirole, on PVIN excitability in layer V M1 motor cortex (Figure 3A). To prevent the influence of spontaneous excitatory and inhibitory inputs on action potential generation, fast glutamatergic and GABAergic transmissions were pharmacologically blocked using DNQX (10 µM) / D-AP5 (50 µM) and picrotoxin (50 µM), respectively. Bath application of quinpirole (2 µM) changed the intrinsic properties of the PVIN sample. A somatic injection of depolarizing current induced more action potentials in the presence of quinpirole for the same injected current, as exemplified in Figure 3A. This was true for all injected currents tested as shown by the frequency/current (F/I) input-output curve (Figure 3B, p < 0.0001, n = 10; 2-way repeated measures ANOVA). Indeed, quinpirole changed the output-input curve of the 10 PVINs tested, shifting it to the left and thus inducing increased excitability. Importantly, the application of the D2R antagonist sulpiride blocked the excitatory effect of quinpirole on PVIN excitability (Figure 3C). Moreover, quinpirole significantly depolarized the PVIN resting potential from −80.19 ± 1.99 mV to −76.44 ± 1.87 mV (p = 0.002; n=10; WSR test), increased their maximal firing frequency (221.1 ± 26.9 Hz to 253.8 ± 28.7 Hz, p = 0.002; WSR test), decreased their rheobase from 235.0 ± 36.6 pA to 180.0 ± 29.1 pA (p = 0.0115; WSR test) and increased their mean input resistance from 127.3 ± 12.9 MΩ to 145.4 ± 18.3 MΩ (p = 0.0020; WSR test) (Figure 3D).

**Figure 3:**
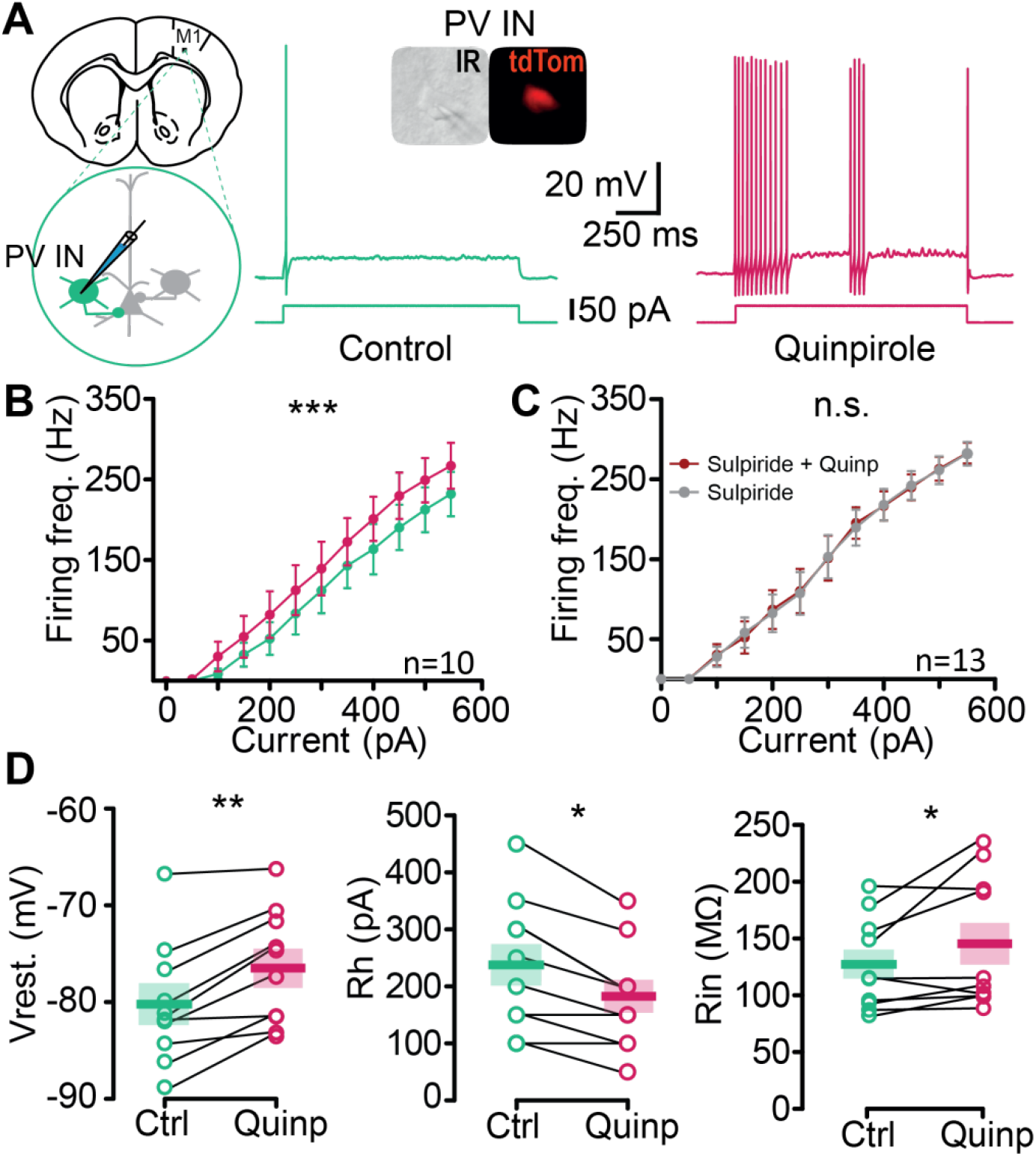
Quinpirole increases the excitability of M1 parvalbumin-expressing interneurons (PVIN). (A) Left, schematic of the experiment. PVINs were identified as tdTomato (tdTom in inset) positive neurons in slices from *PV-Cre:Ai9T* mouse brain. Representative voltage responses to +50 pA current injection in a PVIN in control bath solution (green, middle) and after 10 min of perfusion of the D2R agonist quinpirole (2 µM, red, right). (B) Quinpirole enhanced the firing frequency (Firing freq.) of PVINs and significantly shifted the input-output curve to the left (p < 0.0001, n = 10, F_(1,96)_ = 42.64, 2-way ANOVA). (D) Quinpirole does not increase the firing frequency of PVINs in presence of Sulpiride (p = 0.7645, n = 13, F_(1,144)_ = 0.09011, 2-way ANOVA). Each symbol represents mean ± SEM. (D) Summary of the quinpirole effect on resting membrane potential (Vrest.), maximal firing frequency (Max. freq.), rheobase (Rh) and input resistance (Rin) (WSR test). The thick bar and the color block represent the mean and the SEM, respectively. GABA_A_, NMDA and AMPA/Kainate receptors were blocked throughout all of the recordings with PTX (50 µM), D-AP5 (50 µM) and DNQX (10 µM), respectively.

### D2R activation increases afferent GABAergic synaptic transmission received by PNs

Since PVINs increased their excitability in the presence of quinpirole, we sought to determine whether in the presence of the D2R agonist, individual PNs in layer V received more phasic GABA_A_ receptor-mediated inhibition. We first assessed if quinpirole *per se* changed the intrinsic properties of PNs. On average, a bath application of quinpirole had no effect, neither on the F/I Curve nor on the resting potential or input resistance of the 7 PNs recorded (Figure 4A and Figure 4B). To determine whether PNs received more GABAergic inhibition, we recorded the inhibitory post-synaptic currents (IPSCs) in PNs, i.e. the sIPSCs and mIPSCs that reflect the action potential-dependent and action potential-independent activities of the inhibitory interneuron network, respectively. To specifically study the action of quinpirole on sIPSCs and mIPSCs, and to neutralize the potential confounding influence of excitatory and GABA_B_ neurotransmissions, DNQX (10 µM), D-AP5 (50 µM) and CGP55845 (1 µM) were bath-applied prior to the perfusion of quinpirole. In these conditions (considered as a control condition), robust sIPSCs were observed in all the recorded PNs at a holding potential of −64 mV, confirming GABAergic inhibitory control of PNs by GABAergic interneurons (Figure 4C and 4F).

**Figure 4:**
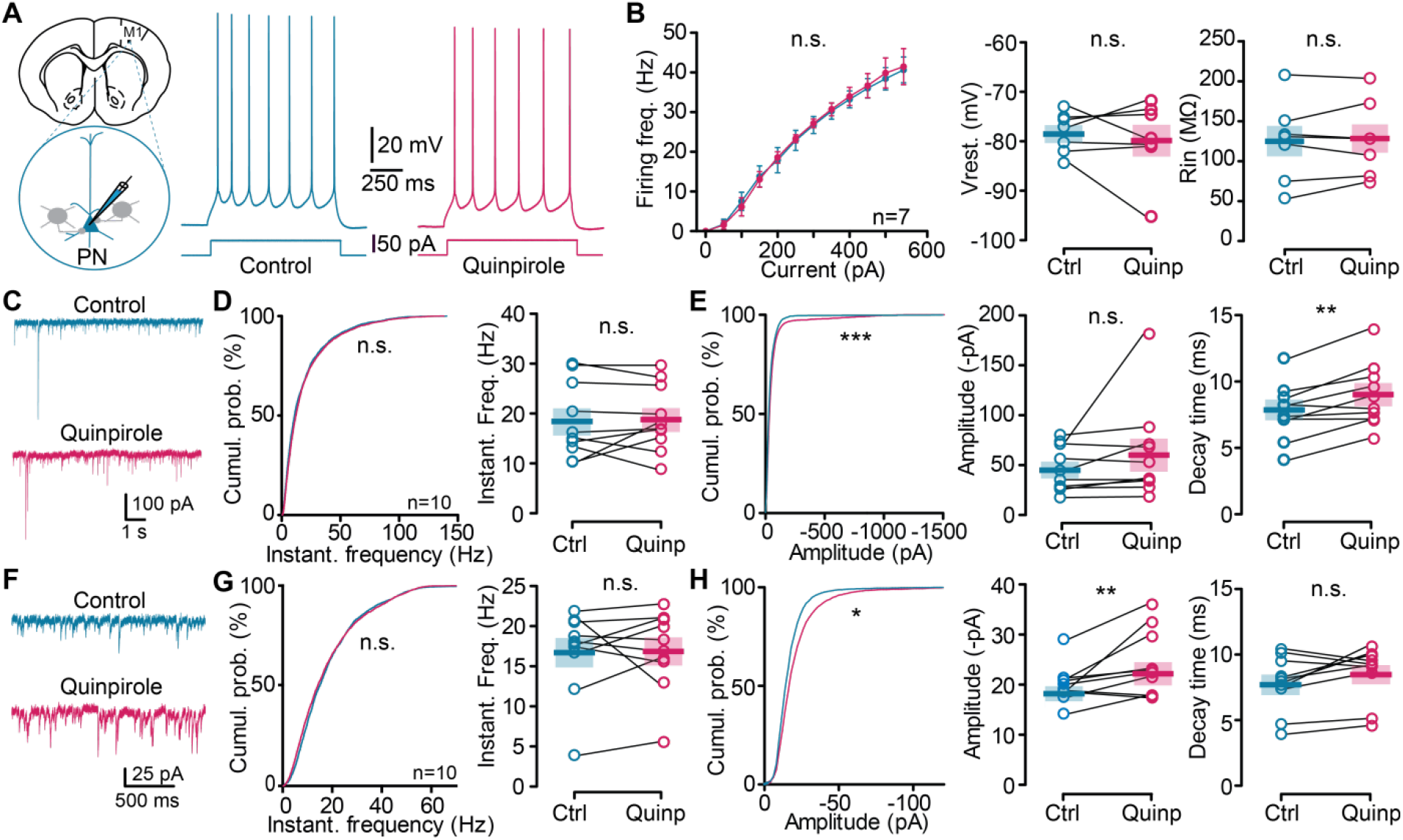
Effect of quinpirole on the electrical activity and spontaneous inhibitory post-synaptic currents (sIPSCs and mIPSCs) of pyramidal neurons. (A) Schematic of the experiment. PNs were identified by their morphology, the absence of tdTomato in their soma as well as their intrinsic properties when possible. Example of voltage responses to +50 pA current injection recorded in a representative PN in control (left, blue) and in quinpirole (right, red). (B) Quinpirole did not change the firing frequency of PNs (p = 0.6453, n=7, F_(1,72)_ = 0.21, 2-way ANOVA), nor the resting potential or input resistance (p = 1.000 and p = 0.8982, respectively; n = 7, WSR test). (C) Representative traces of sIPSCs recorded from a M1 PN (left) in control conditions (top trace, blue) and in the presence of quinpirole (bottom trace, red). (D) Cumulative distribution (left) and mean (right) of sIPSC instantaneous frequency in the control (blue) and in the presence of quinpirole (red). No differences were observed (p = 0.9987, Kolmogorov-Smirnov and p > 0.9999, WSR test). (E) Cumulative distribution (left) and mean (middle) of sIPSC amplitude and value of tau (right). Note that cumulative distribution of the amplitude and decay time differed significantly between control and quinpirole conditions (p < 0.0001, Kolmogorov-Smirnov, and p = 0.0059, WSR test, n = 10). (F-H) Same representations as in C, D, E for mIPSCs. Note that similarly to sIPSCs, only the cumulative distribution (p < 0.05, K-S test) and mean of mIPSC amplitude differed significantly between control and quinpirole conditions (p = 0.0098; n = 10, WSR test).

The effects of 2 μM quinpirole on sIPSCs were studied on 10 neurons. Quinpirole increased the amplitude (Figure 4E) without changing the frequency of the sIPSCs (Figure 4D). Indeed, the cumulative probabilities of the frequency of sIPSCs in the control and the quinpirole groups were similar (Figure 4D, P >0.05, K-S test). However, the cumulative probability of the amplitude of the sIPSCs showed an increase in the quinpirole group (Figure 4E, p < 0.0001, K-S test) compared to the control group. Moreover, quinpirole significantly increased the decay time from 7.68 ms ± 0.67 to 9.07 ± 0.75 ms (p = 0.059, n = 10; WSR test). Next, we examined the effect of quinpirole in PNs in the presence of 1 µm TTX, to isolate mIPSCs (Figure 4F). As for sIPSCs, analysis of the cumulative probability (Figure 4G and 4H) with the K-S test revealed that D2R activation increased mIPSC amplitude with no effect on their frequency. The decay time of IPSC was significantly increased from 7.66 ± 0.65 to 8.88 ± 0.70 ms (p = 0.0273, n = 10; WSR test). The frequency was unchanged on average, but it is important to note that quinpirole had a variable effect on individual neurons.

### D2R activation enhances GABAergic transmission at PVIN-PN synapses

Our results on GABAergic IPSCs suggested that D2R activation by quinpirole induced more activity in the inhibitory network. However, the increase observed may be due to any type of inhibitory IN. To determine whether quinpirole changes synaptic transmission between PVINs and PNs, we used optogenetics to selectively study PVIN-PN synapse properties (Figure 5). We expressed the channelrhodopsin ChR2 in PVINs *via* local viral transfection in M1 of PV-Cre:Ai9T mice using an AAV2.5-EF1a-DIO-hChR2(H134R)-EYFP vector (Figure 5A). We used 473 nm light flashes to stimulate PVINs while recording from PNs. We first confirmed that 1 ms flashes of light were able to reliably trigger action potentials in PVINs. As illustrated by the raster plot in Figure 5B, each flash in the train evoked one or two action potentials in the transfected PVIN. In a second step, we recorded the optically-evoked IPSCs from PNs (Figure 5C). PNs were identified as described previously (Figure 4) and displayed a PN-typical firing pattern upon depolarizing current steps (Figure 5C). Light flashes reliably elicited evoked inhibitory post-synaptic currents (eIPSCs) in PNs, which were potentiated by bath application of 2 μM quinpirole (Figure 5D), increasing their mean amplitude from 280.3 ± 68.52 pA to 321.6 ± 75.67 pA (p = 0.0371, n = 10; WSR test). This result strongly suggested that GABAergic synaptic transmission between PVINs and PNs was enhanced by quinpirole. We further characterized the short-term plasticity of the PVINs-PNs synapses using 10 flashes of 1 ms at 10 Hz; Figure 5E). The inhibitory inputs to PNs showed pronounced synaptic short-term depression, but bath-applied quinpirole did not change the profile of synaptic transmission, which remained depressed (Figure 5F).

**Figure 5:**
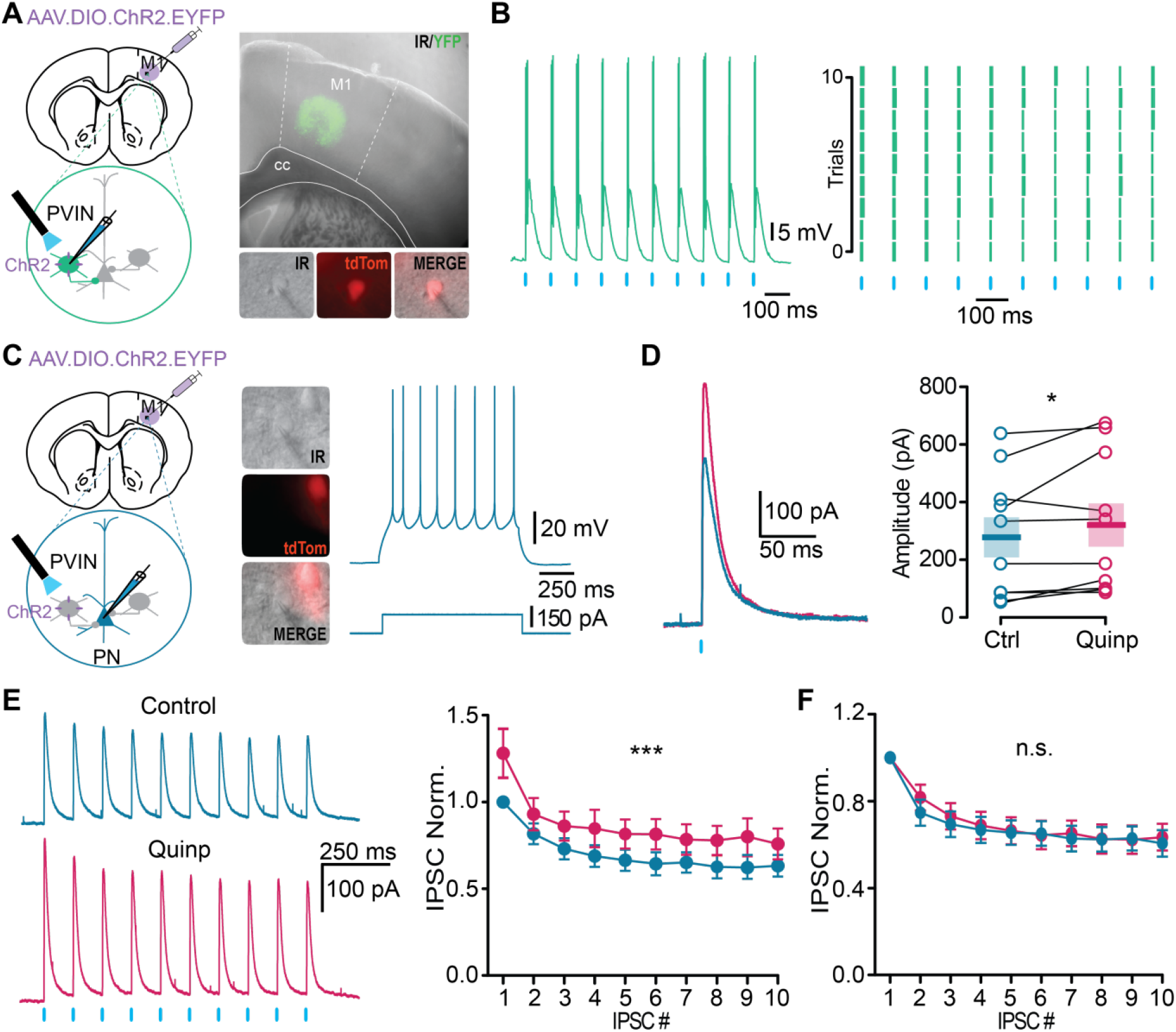
Dopamine increases GABAergic synaptic transmission at the PVIN-PN synapse. (A) Schematic of the experiment. An AAV.DIO.ChR2.EYFP virus was injected in M1 2 weeks before *ex vivo* recordings. Representative slice showing the expression of ChR2-EYFP in M1. PVINs (tdTom-positive) were patched to verify our ability to manipulate their activity. (B) Light reliably induced action potentials in a PVIN (left). Each flash of light (blue line) evoked 1 or 2 action potentials in PVINs as presented in an example (left) of an intracellularly recorded PVIN and in the raster plot of spiking in different trials (right). Each blue tick represents a flash of light (473 nm, 1 ms) and each green tick represents a spike. (C) Schematic of the recording configuration from a post-synaptic PN during photoactivation of PVINs (left). Soma of PNs had a triangular shape, were tdTom-negative and PV-negative (middle). Representative firing pattern of recorded PNs to a 150 pA, 500 ms current step (blue, right). (D) Sample traces and quantification of light-evoked IPSCs recorded in the same PN before (blue, control trace) and after 10 min of quinpirole perfusion (red). Mean of the amplitude of the evoked response (p = 0.0371, n = 10, WSR). Mean and SEM are represented. (E) Sample traces and quantification of responses to repetitive photostimulation (10 Hz) recorded before (blue) and after bath application of quinpirole (red). Photoactivation of PVINs produced large initial IPSCs that depress rapidly. In the graph, the IPSC amplitudes were normalized to that of the first IPSC in the control condition for each neuron recorded (p < 0.0001, n = 10, F(_1,180_) = 19.36). (F) Short-term synaptic dynamics of the evoked IPSCs in PNs induced by the photoactivation of PVINs were not changed in presence of quinpirole. IPSC amplitudes were normalized to the first IPSC of the train in each condition (p = 0.1563, n = 10, F(_1,180_) = 0.4749).

## Discussion

In the present study, we first performed quantitative mapping of M1 neuronal populations expressing D2R. These neurons are largely present in layers II/III and layer V, and are mainly PVINs in layer V, based on immunochemistry and electrophysiological characterization. Then, combining electrophysiology and optogenetics, we demonstrated *ex vivo* that the activation of D2R robustly increases the excitability of PVINs and enhances the synaptic transmission between PVINs and PNs.

### D2R expressing cells in layer V of M1

Previous studies have shown that M1 cortical neurons express both D1 and D2 classes of DA receptors (Lidow et al., 1989; Seamans and Yang, 2004) and receive direct DA projections from VTA and SNc *via* meso-cortical pathways (Descarries et al., 1987). For many years, cortical D2R has been a focus of interest because of its involvement in many cognitive functions initiated or modulated by DA. However, the relatively low expression of cortical D2R makes its detection very difficult. Consequently, it is more difficult to identify the nature of the neurons expressing D2R, a difficulty further amplified by the massive heterogeneity of neurons. Most studies of cortical D2R have focused on the prefrontal cortex and several studies have detected the presence of D2R mRNA in PFC by *in situ* hybridization, revealing its expression in pyramidal neurons and minor subtypes of interneurons (Gaspar et al., 1995). Recently, technical limitations were overcome with a highly sensitive and multimodal approach to map cortical D2R-expressing neurons (Drd2-Cre:Ribotag mouse), which has allowed the identification of previously uncharacterized clusters of D2R-expressing neurons in limbic and sensory regions of the adult mouse brain (Khlghatyan et al., 2018). Unfortunately, the authors did not perform quantitative mapping of the M1 neuronal populations expressing D2R. In the present study, using both the Drd2-Cre:Ribotag and Drd2-Cre:Ai9T mouse lines, we show that D2R-expressing cells are distributed in all cortical layers of M1 and broadly expressed in layer V. The molecular characterization of D2R-expressing cells in layer V revealed a majority of PVINs and to a lesser extent, populations of CB- and NPY-positive cells. Electrophysiological characterization in the Drd2-Cre:Ai9T mouse line revealed 3 main classes of neurons expressing D2R in layer V of M1: FS, RSNP and PN. The majority of the D2R-expressing cells are FS neurons, which are also mainly PV-positive neurons. Since PVINs account for a quarter of the D2R-positive neurons, they are likely to play a specific role, still unknown, as a target for the DA modulation of cortical microcircuits in M1.

### D2R modulation of intrinsic excitability of PVINs in M1

Although dopaminergic fibers and DA receptors in M1 have been clearly demonstrated (Descarries et al., 1987; Hosp et al., 2011), their functional significance remains poorly understood. Conflicting evidence indicates excitatory and inhibitory effects on electrical activity *in vivo* (Vitrac and Benoit-Marand, 2017). Most studies have not tested specific cell types and there is no data available regarding DA modulation of specific subpopulations of interneurons in M1. Here, we examined PVINs of layer V and found that activating D2R caused their depolarization and an increase in their excitability. Such an excitatory effect of the D2R agonist quinpirole on interneurons has been already observed on interneurons in prefrontal cortex (PFC) slices from adult mice (Tseng and O’Donnell, 2006). As cell excitability was determined by assessing the response to intracellular injection alone and in the presence of fast synaptic blockers, this is likely to reflect the post-synaptic effects of the agonist and not a modulation of pre-synaptic afferents or neurons. However, further studies will be required to determine whether this excitatory effect reflects a direct D2R post-synaptic action on PVINs as observed in PFC (Tseng, 2004), or activation of D2R autoreceptors and release of the co-transmitter neurotensin, which is present in a subpopulation of DA neurons from the VTA projecting to PFC (Petrie, 2005) that may also project to M1.

### D2R modulation of GABAergic synaptic transmission in M1

Proper brain function depends on a correct balance between excitatory and inhibitory signaling (Markram et al., 2015) and PVINs are crucial for such network functionality. Indeed, they exert powerful actions on cortical network activity by contributing to feedback and feed-forward inhibition of pyramidal neurons (Hu et al., 2014). To determine if the D2R agonist quinpirole modulates the afferent GABAergic synaptic transmission to PN in M1, we examined the IPSCs in PNs. Measurement of the changes in sIPSCs and mIPSCs are a sensitive means to estimate the locus of a drug effect. Typically, changes in IPSC amplitudes are associated with a post-synaptic site of modulator action, whereas changes in IPSC frequency are likely to be due to an interaction with a presynaptic site that changes the probability of transmitter release (Lupica, 1995). Since the excitability of PVINs was increased by quinpirole, we expected an increase in IPSC frequency, but this was not the case. Here, quinpirole increased the amplitude of both sIPSCs and mIPSCs recorded in PNs with no effect on their frequency. One possible explanation is that since the PVINs are not spontaneously firing at their resting potential, the 5 mV depolarization generated by quinpirole may not be large enough to raise the resting membrane potential to the spike threshold in the absence of excitatory transmission. Another possible explanation is the various origins of the GABAergic IPSCs. We studied all the GABAergic inhibitory currents received by PNs and cannot exclude an effect of quinpirole on other GABAergic INs that may mask the effect on frequency. Finally, this may occur only in a subset of PVIN-PN synapses.

### D2R modulation of PVINs-PN synaptic transmission in M1

In order to be specific to PVIN-PNs synapses and to overcome the fact that these PVINs do not spontaneously fire in slices, we used optogenetics. Our results show that bath application of quinpirole potentiated the optically-evoked eIPSCs in PNs. Using a 10 Hz train of stimulations, we showed an adaptive depression of this synapse from the second optical stimulation that persisted with quinpirole. These observations suggest that quinpirole mainly acts on post-synaptic sites and show that the adaptive depression is maintained. It is also possible that quinpirole-induced depolarization of PVIN membrane potential allows the recruitment of a greater number of neurons during light activation, which can account for the increase in optically-evoked IPSC amplitude without changes in the depression profile. Finally, bath application of quinpirole clearly modulates the intrinsic properties of PVINs. However, as the timing of DA release is critical for plasticity induction (Yagishita et al., 2014), investigating how endogenous DA release controls PVIN-PN GABAergic synaptic transmission and plasticity using optogenetics is crucial.

### Functional implications

M1 is particularly important in acquisition and maintenance of motor skills and is a central locus for motor learning. Indeed, pharmacological or optogenetic inactivation of M1 is highly effective in reducing motor aptitude (Peters et al., 2014; Guo et al., 2015; Otchy et al., 2015). The lesion of M1 before training abolishes the ability to learn stereotyped movements but does not impair the execution of an already learned motor skill (Guo et al., 2015; Kawai et al., 2015), demonstrating a role for M1 in ‘tutoring’ subcortical circuits during skill learning. Moreover, recent studies have shown that DA plays a key role in motor learning and memory in M1 (Molina-Luna et al., 2009; Leemburg et al., 2018), particularly in spine regulation and synaptic plasticity (Xu et al., 2009; Guo et al., 2015). Interestingly, it has been recently shown that PVINs exhibit a gradual increase in axonal boutons during motor training (Chen et al., 2015). As we show that the activity of PVINs can be modulated by activation of D2R in M1, these data suggest that PVINs and D2R may be crucial for learning sophisticated motor sequences. Interestingly, it has been shown that striatal PVINs enhance behavioral performance in a reward-conditioning task, but their contribution declines as learning progresses (Lee et al., 2017), suggesting dynamic involvement during the learning of the task. Thus, we expect that following the loss of DA in M1 in conditions such as Parkinson’s disease, plasticity of PVINs in M1 will be altered and can lead to the cognitive deficits observed in this pathology.

## Contributions

JC, LL, EV, AT, LDZ, JB and MLBJ performed the experiments; JC, LL, EV, AT and MLBJ analyzed the data; EV, AT, JB and MLBJ wrote the manuscript.

## Conflict of interest statement

The authors declare no competing financial interests.

## Acknowledgments

This work was supported by CNRS, University of Bordeaux, Inserm, Fondation pour la Recherche Médicale (FRM) and the French National Research Agency ANR-DOPAFEAR (to EV). JC received a Ph.D fellowship from the FRM (ECO201806006853 – Fondation Yolande Calvet).

We thank G. Dabee for animal care and M. Goillandeau for assistance with his software for the detection of spontaneous and mini IPSCs. We are also grateful to Dr. Patricia Gongal (Innovology, Canada) for language assistance on the manuscript.

